# Structures of a lipin/Pah phosphatidic acid phosphatase in distinct catalytic states reveal a signature motif for substrate recognition

**DOI:** 10.1101/2025.06.16.660012

**Authors:** Tereza Vitkovska, Franceine S. Welcome, Valerie I. Khayyo, Shujuan Gao, Troy Wymore, Michael V. Airola

## Abstract

Lipin/Pah phosphatidic acid phosphatases (PAPs) are Mg^2+^-dependent enzymes that catalyze the dephosphorylation of phosphatidic acid (PA) to produce diacylglycerol. Deficiency of lipin PAP activity in humans results in inflammatory disorders such as rhabdomyolysis and Majeed syndrome. Previously, we reported the first PAP enzyme structures of *Tetrahymena thermophila* (*Tt)* Pah2 at 3.0Å resolution. Here, we present five new higher resolution (1.95-2.40Å) structures of *Tt* Pah2, in distinct catalytic states, which represent active states of catalysis, including a phospho-Asp transition state mimic, and an inactive state with a distorted active site. The structures, in conjunction with flexible docking simulations and biochemical analysis, connect two highly conserved aspartate and arginine residues in magnesium coordination and recognition of the substrate PA. Overall, this provides a structural basis for catalysis and defines a signature Asp-Arg motif in lipin/Pah PAPs that enables recognition of their lipid substrate PA, providing insight into how the HAD domain of lipin/Pah PAPs evolved to act on a membrane embedded substrate.

## INTRODUCTION

Lipin/Pah phosphatidic acid phosphatases (PAPs) are magnesium-dependent enzymes that are evolutionarily conserved from yeast to humans^1,2^. They catalyze the penultimate step of triacylglycerol (TAG) synthesis, through the conversion of phosphatidic acid (PA) to diacylglycerol (DAG)^3^ (**Fig. 1**). Since PA and DAG are precursors for various phospholipids, PAP enzymes also regulate phospholipid biosynthesis^4,5^ (**Fig. 1**). There are three lipin isoforms with multiple splicing variants in humans and mice^6,7^, one PA phosphohydrolase (Pah) in the yeast *Saccharomyces cerevisiae* (*Sc*)^1^, and two Pah isoforms (Pah1 and Pah2) in the ciliate *Tetrahymena thermophila* (*Tt*)^8^.

**Figure 1.**
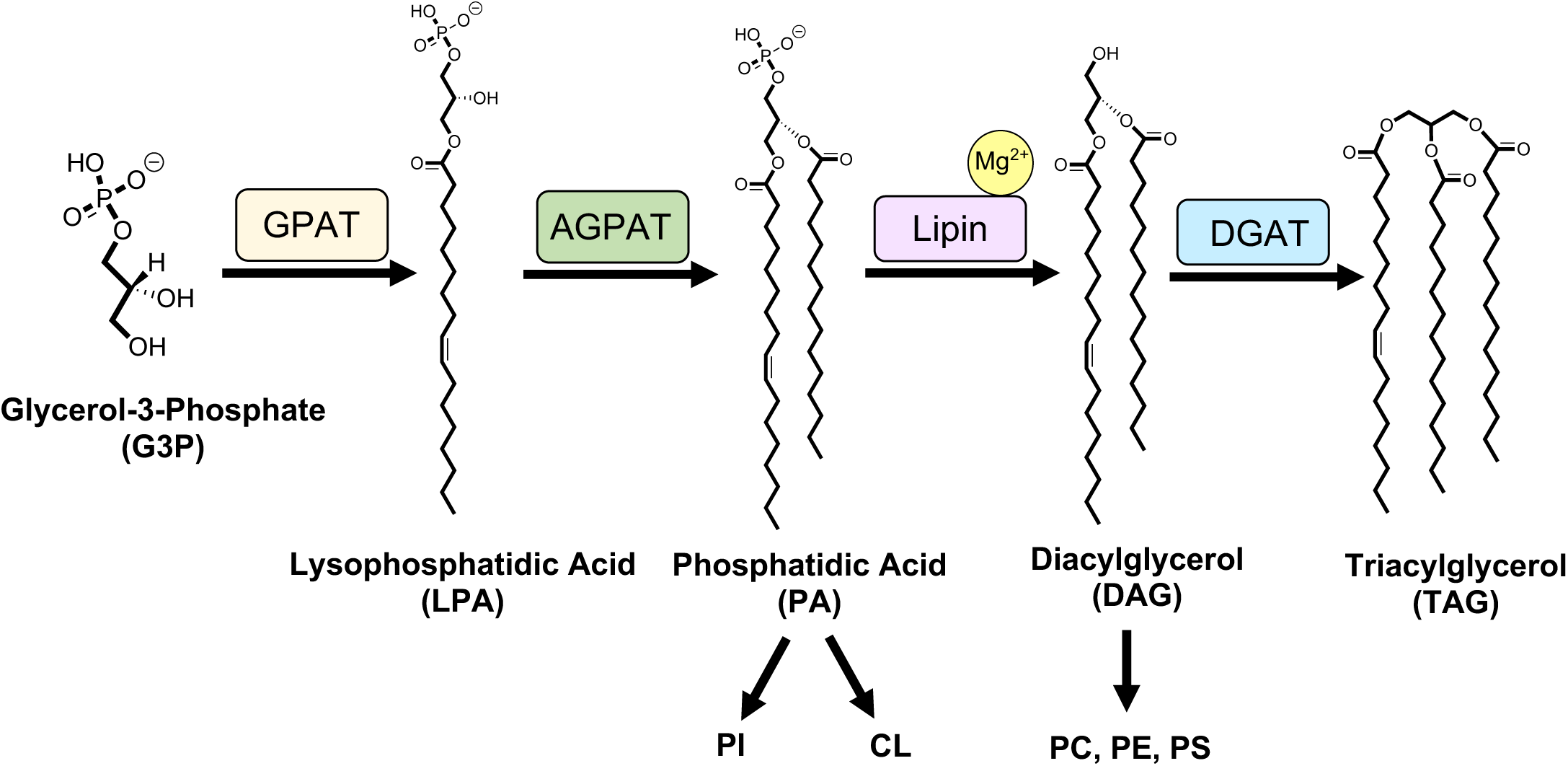
Glycerol-3-phosphate pathway with structures of intermediates. Lipin is a magnesium dependent phosphatidic acid phosphatase (PAP) that catalyzes the penultimate step of the glycerol-3-phosphate pathway. This reaction is essential for generation of the primary energy storage molecule in eukaryotes, triacylglycerol (TAG), and major membrane phospholipids phosphatidylcholine (PC), phosphatidylethanolamine (PE) and phosphatidylserine (PS). Lipin preferentially hydrolyzes PA over other intermediates of this pathway, like lysophosphatidic acid (LPA). Glycerol-3-phosphate is acylated to generate lysophosphatidic acid (LPA) by glycerol phosphate acyltransferase (GPAT). LPA is further acylated to generate phosphatidic acid (PA) by acylglycerol phosphate acyltransferase (AGPAT). Lipin dephosphorylates PA to generate diacylglycerol (DAG). In the final step, diacylglycerol acyltransferase (DGAT), acylates DAG to generate TAG.

The importance of PAP enzymes is exemplified by the various diseases in humans, and phenotypes in mice and yeast that result from deficiency in lipin or Pah PAP activity. In humans, lipin-1 PAP loss-of-function mutations are associated with Rhabdomyolysis, a condition characterized by fever and muscle tissue degradation, which causes renal failure^9–11^. Majeed syndrome, seen in people with lipin-2 inactivating point mutations, is a painful inflammatory disorder, episodes of which cause anemia, fever, and pain in bones and joints^12–15^. Phenotypes in mice are associated with fatty liver dystrophy^7^ and neuropathy^16^, while phenotypes in the yeast *S. cerevisiae* include sensitivity to both heat and cold, defects in using non-fermentable carbon sources, and nuclear/ER membrane expansion^17,18^. Thus, lipin/Pah PAP biology has metabolic implications.

Structurally, all lipin/Pah PAPs contain highly conserved N-Lip and C-Lip regions that combine to form two domains: an immunoglobulin-like (Ig-like) domain and a HAD-like catalytic domain^19^. The Ig-like domain is formed through co-folding of the N-Lip region with a small portion of the C-Lip^19^. The remainder of the C-Lip adopts a Rossmann-fold, forming the haloacid dehalogenase-like (HAD-like) domain and harbors a conserved catalytic motif DxDxT^20–22^, where the first Asp coordinates a Mg2+ ion that increases its nucleophilicity, and the second Asp participates in acid/base catalysis (**Fig. 2**). Beyond the N-Lip and C-Lip, PAP architecture can differ. Mammalian, plant, and some ciliate lipin/Pah PAPs contain a dimerization and membrane binding middle lipin (M-Lip) domain^23^, while *Sc* Pah1 contains an RP domain between the N-Lip and C-Lip regions that regulates Pah1 phosphorylation^24^. Despite the middle domain variations, the N-Lip and C-Lip regions are sufficient for catalytic function in several PAP enzymes. This includes *Tt* Pah2, which endogenously fuses these regions^8,19^, mouse lipin-2^19^, human lipin-1^25^, and *Sc* Pah1^25,26^. Consistently, all known disease and phenotype-associated mutations in human and mice lipins are found within the N-Lip or C-Lip regions^7,9–13,15,16,27–30^.

**Figure 2.**
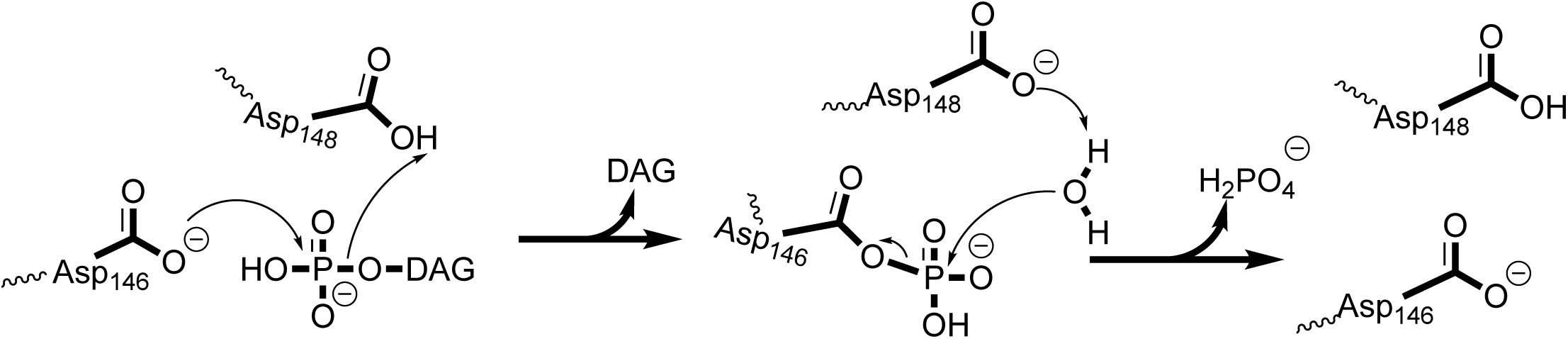
Proposed mechanism of catalysis of magnesium dependent lipin/Pah PAPs. Lipin/Pah PAPs harbor a DxDxT catalytic motif characteristic of the haloacid dehalogenase superfamily (HADSF). The first catalytic Asp (Asp146) attacks the slightly positively charged phosphorus of phosphatidic acid (PA) forming the trigonal bipyramidal phospho-Asp intermediate, while the DAG leaving group pulls a hydrogen from the second Asp (Asp148), leaving the active site in the first step. In the next step, Asp148 can then activate a water molecule that attacks the phospho-Asp intermediate, freeing the phosphate, regenerating the original state of the enzyme.

The catalytic domain of lipin/Pah PAPs belongs to the widespread haloacid dehalogenase (HAD) superfamily of enzymes, which mainly act as phosphatases on a variety of substrate types (e.g. phospholipids, phospho-proteins, small molecules)^20–22^. The substrate specificity in HAD phosphatases is often dictated by cap domains that sit above or adjacent to the active site and make direct interactions with substrates^20,31^. HAD phosphatases are classified as either type 0 (no cap), type 1 (domain in cap region 1), or type 2 (domain in cap region 2), dependent on the absence or presence of a cap domain, and which secondary structural elements of the Rossmann-like fold the cap domain is located between^20,22,32^.

*Tt* Pah2 is the only HAD phosphatase with an experimental structure that acts on a membrane embedded substrate. Our previous structures revealed that *Tt* Pah2 and other Mg^2+^ dependent PAPs lack a canonical cap ‘domain’ but have relatively short ∼20 residue peptides inserted into both cap region 1 and cap region 2^19^. Both of these cap regions were either disordered in prior *Tt* Pah2 structures or found in differing conformations, thus reflecting their flexibility. Overall, this suggested that lipin/Pah PAPs might interact with the lipid substrate, PA, through either cap region 1 or 2, most likely through the glycerol backbone of PA, which would be consistent with the lack of acyl-chain specificity in both Sc Pah1 and lipin PAPs^2^.

Here, we sought to determine the structural basis for lipin/Pah PAPs catalysis and substrate recognition of PA. Towards this goal, we determined new high-resolution structures of *Tt* Pah2 either in an inactive state or in different active Mg^2+^ bound states. These structures revealed additional conformational flexibility of the peptides in both cap regions, and visualized a tungstate bound state that mimicked a transition state of catalysis. Point mutations, coupled with biochemical characterization, identified an Asp and Arg residue critical for both PA catalysis and recognition. These Asp and Arg residues were highly conserved in PAP enzymes but are distinct from the canonical and universally conserved residues in HAD phosphatases with known roles in catalysis^20,22^. Consistent with a role in PA recognition, flexible docking analysis identified interaction with the glycerol backbone of PA by the Arg residue. Overall, this provides new insight into the structural basis of substrate recognition and catalysis by lipin/Pah PAP enzymes.

## RESULTS

### Structures of *Tt* Pah2 in distinct conformations

*Tt* Pah2 represents the only magnesium dependent phosphatidic acid phosphatase (PAP) enzyme with available experimental structures. Previously, we determined crystal structures of *Tt* Pah2 bound to either Mg^2+^ or Ca^2+^ at 3.0 Å resolution^19^. To determine higher resolution structures and gain insight into the structural mechanisms of lipin/Pah PAP catalysis and PA recognition, we generated constructs of *Tt* Pah2 that lacked the N-terminal amphipathic helix (Δhelix), which was disordered in prior structures due to high flexibility^19^. Our strategy yielded four new Δhelix *Tt* Pah2 structures and one additional structure that retained the N-terminal helix **(Fig. 3**, **Table 1)** that were refined to resolutions of 1.95-2.4Å. The five structures belonged to three unique space groups that visualized *Tt* Pah2 in either distinct conformations or stages of catalysis.

**Figure 3.**
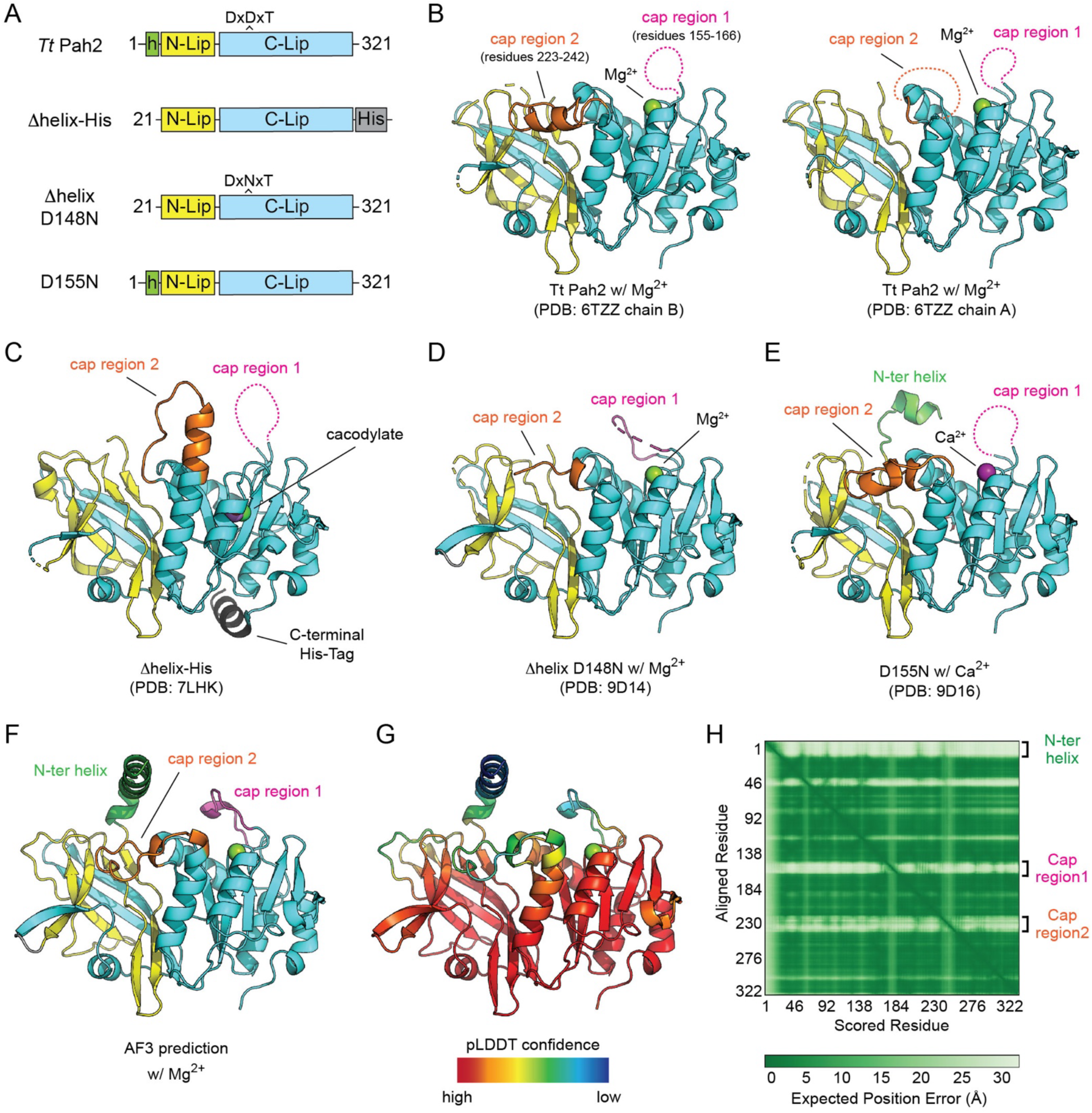
Tt Pah2 crystal structures reveal conformational flexibility of cap regions. **(A)** Domain architectures of the different crystallized fragments and point mutants of *Tt* Pah2. **(B)** Cartoon representation of *Tt* Pah2 crystallized with magnesium^19^. Cap region 1 (magenta) is disordered in both molecules (chain A and B) of the asymmetric unit, while Cap region 2 (orange) forms a helix in chain B but is disordered in chain A. **(C)** Cartoon representation of *Tt* Pah2 Δhelix-His that highlights cap region 2 forming a continuous helix and a disordered cap region 1. **(D)** Cartoon representation of the *Tt* Pah2 Δhelix D148N with magnesium (green sphere) that highlights the partially ordered cap regions 1 and 2. **(E)** Cartoon representation of *Tt* Pah2 D155N that highlights the helical cap region 2, the disordered cap region 1, and the partially ordered N-terminal helix (green). **(F)** AlphaFold 3 structure prediction of *Tt* Pah2 with a magnesium ion (green) bound, showing partially ordered peptides in cap region 1 and cap region 2. **(G)** Corresponding pLDDT confidence values for predicted structure of *Tt* Pah2, showing low confidence in structure of N-terminal amphipathic helix and cap regions 1 and cap region 2. **(H)** Predicated Alignment error (PAE) plot showing the expected position error for predicted structure of *Tt* Pah2 showing peptides in both cap regions and the N-terminal amphipathic helix have low confidence in their position relative to the rest of the protein.

**Table 1.**
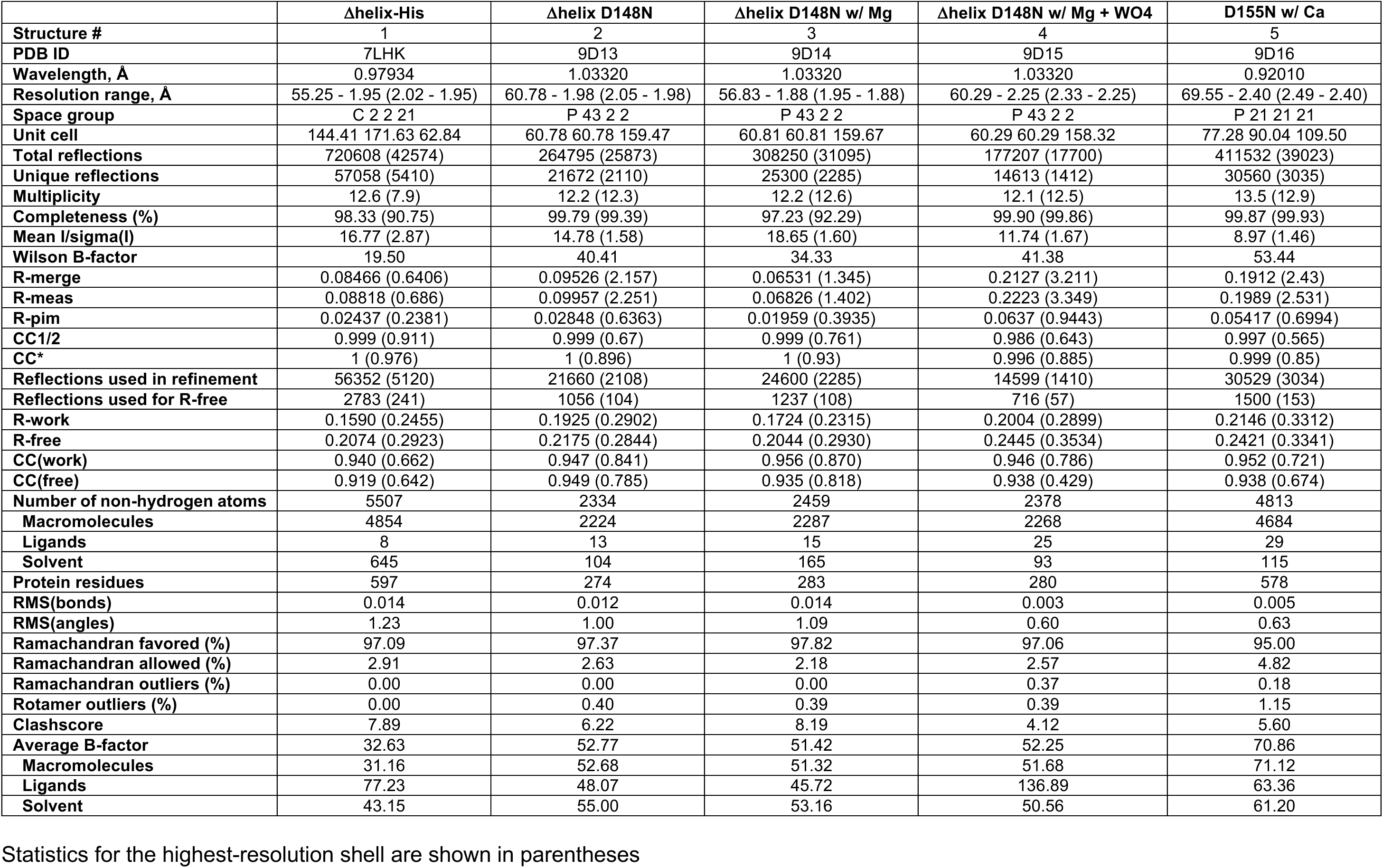
Data Collection and Refinement Statistics.

Structure 1 was determined using a Δhelix-His construct that lacked the N-terminal helix but included a C-terminal His-tag **(Fig. 3C)**. The Δhelix-His structure represented an inactive state due to covalent modification of Cys276 by cacodylate, which was present in the crystallization solution. Cacodylate modification of Cys276, which lies below the nucleophilic Asp of the DxDxT motif, induced a major rearrangement of active site loops and residues, including an Asn residue (N268) that binds Mg^2+^, the adjacent Arg residue (R269) we implicate in PA recognition below, and an Asp residue (D272) that is mutated to His or Asn in rhabdomyolysis^33,34^, which formed a hydrogen bond with a Mg^2+^ bound water in other active states of catalysis.

Structures 2, 3, and 4 were determined using a Δhelix construct with a D148N point mutant that modified the DxDxT motif to be DxNxT **(Fig. 3A, 3D)**. Mutating polar residues in the active site to nonpolar equivalent residues has been demonstrated to stabilize HAD phosphatases, enhancing structure resolution^35,36^. The Δhelix D148N point mutant was crystallized in an apo state, Mg^2+^-bound, and Mg^2+^-tungstate bound states by crystal soaking. This allowed visualization of conformational changes that may occur during different stages of catalysis (discussed below). Structure 5 was determined using a construct that retained the N-terminal helix but contained a D155N point mutant. Crystal packing involved interactions of the N-terminal helix with another molecule, with the N-terminal helix partially ordered in this structure **(Fig. 3E)**.

### The two membrane facing cap peptides in *Tt* Pah2 are flexible and adopt variable conformations

Substrate specificity in HAD phosphatases is typically dictated by cap domains that are inserted between secondary structural elements of the Rossmann-fold of the HAD core^31,32^. Dependent on the absence or presence of a cap domain, and which secondary structural elements the cap domain is located between, HAD phosphatases can be classified as either type 0, type 1, or type 2^20,32^. *Tt* Pah2 and other Mg^2+^ dependent PAPs have relatively short 20 residues peptides inserted into both cap region 1 and cap region 2, but these peptides do not form domains.

Comparison of the five new *Tt* Pah2 crystal structures, with our previous two structures, identified significant conformational flexibility in both cap region 1, which lies just above the DxDxT active site motif, against the membrane interface, and cap region 2, which is parallel to the membrane interface and resides near the active site between the HAD domain and Immunoglobulin-like (Ig-like) domain of *Tt* Pah2 (**Fig. 3B-E**). Notably, these cap regions were disordered in the structures, unless they were involved in crystal contacts that trapped them in a single conformation. Consistently, alphafold3 predictions **(Fig. 3F)** suggested these were also the most flexible regions based on the comparatively lower predicted local distance difference test (pLDDT) **(Fig. 3G)** and predicted alignment error **(Fig. 3H)** scores.

Cap region 1 was disordered, completely lacking electron density, in all prior and some new structures. The exceptions were structures 2, 3, and 4 determined using the Δhelix D148N construct. These structures had a partially ordered cap region 1 that lacked secondary structure and had higher B-factors than the residues in the HAD core, reflecting their relative flexibility. This suggests that cap region 1 is a dynamic region that can adopt multiple conformations, which could be influenced by either membrane or substrate binding.

A comparative structural analysis of Cap region 2 revealed more dramatically different conformations. In prior structures, cap region 2 was either completely disordered or formed an alpha helix that ran parallel to the membrane binding interface (**Fig. 3B**). In structures 2, 3, 4, and 5, the relative parallel orientation to the membrane was retained but the secondary structure and exact positioning was varied. In sharp contrast, in the Δhelix-His structure (Structure 1), cap region 2 formed an upward facing alpha-helix that was continuous with the prior helix and would be perpendicular to the membrane interface (**Fig. 3C**). As mentioned above, this unique conformation was formed through crystal contacts, with the continuous alpha helix packing into the active site of an adjacent molecule.

### Structural insights into magnesium binding and transition state stabilization

The Δhelix D148N structures were crystallized without any cofactors and then soaked with either magnesium, or magnesium and tungstate to generate three structures with active sites in distinct states. Comparison of the apo and magnesium bound structure saw replacement of a water molecule with a hexa-coordinated magnesium ion in the active site, characteristic of a HAD phosphatase (**Fig. 4A, B)**. As expected, three of the magnesium coordination sites were occupied by the sidechain of the catalytic Asp146, the sidechain of Asn268, and the carbonyl oxygen of the D148N main chain **(Fig. 4B)**. Two water molecules occupied the 4^th^ and 5^th^ coordination sites, with one of the water molecules stabilized by a hydrogen bond with Asp272 **(Fig. 4B)**. Notably, two separate point mutations in the analogous Asp residue in human lipin 1α (D804N/H) were recently associated with rhabdomyolysis, thus the role of Asp272 in *Tt* Pah2 provides a structural rationale for this disease associated mutation in humans.

**Figure 4.**
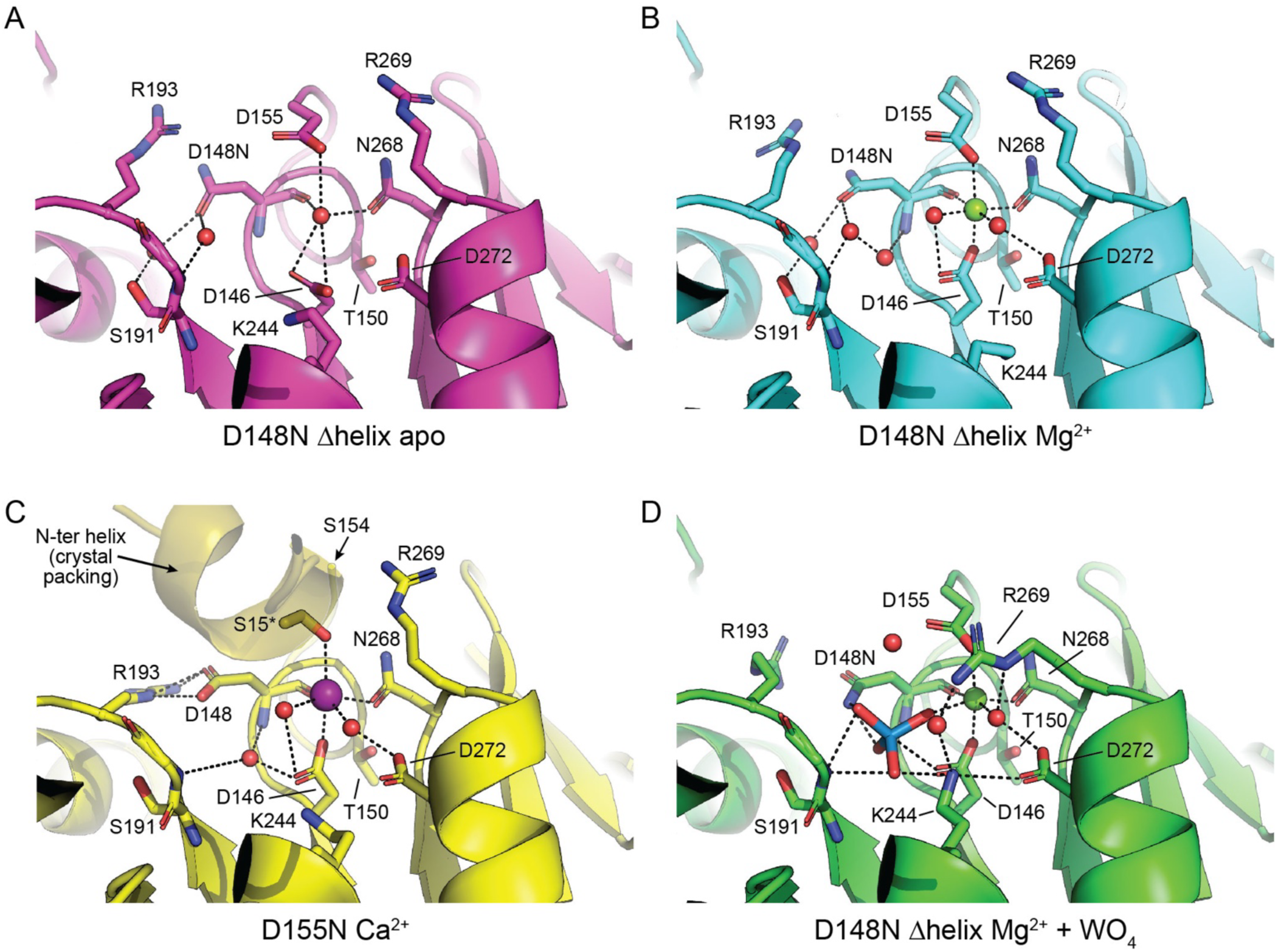
Active sites of Tt Pah2 in different catalytic states. **(A-B)** *Tt* Pah2 active sites from the D148N Δhelix structures in (A) apo and (B) magnesium and bound states. **(C)** *Tt* Pah2 active site from the D155N structure with bound calcium. **(D)** *Tt* Pah2 active site from the D148N Δhelix structure in the magnesium and tungstate bound state. D155 occupies the distal coordination site of magnesium. A conformational change in Lys244 and Arg269 is observed upon tungstate binding. Residue 155 becomes disordered upon substitution to Asn (D155N) with Ser15 from the N-terminal helix of another molecule in the asymmetric unit occupying the distal calcium coordination site. Water molecules are shown as red spheres. Magnesium and calcium ions are shown as green and purple spheres, respectively. Dashed lines represent hydrogen bonds, salt bridges, and polar contacts.

The sixth and final magnesium coordination site was occupied by the carboxylic acid group from the sidechain of Asp155 and was located distal to the catalytic Asp146 on the opposite side of the magnesium ion **(Fig. 4B)**. In the apo structure, Asp155 was observed in a similar conformation but coordinated the water molecule in place of the magnesium **(Fig. 4A)**. We note that there was no electron density for Asp155 in our prior structures determined with either magnesium or calcium, which we assume was due to the low resolution of these structures.

Other HAD phosphatases have similar magnesium coordination^20–22,32,37,38^, however we sought to increase our confidence in our modeling by determining the structure of a D155N point mutant. This structure (Structure 5) was determined in the presence of calcium. Notably, there was no observable electron density for the entire D155N residue, and the sixth coordination site of calcium was now occupied by a serine residue from the N-terminal helix from an adjacent molecule in the asymmetric unit **(Fig. 4D)**. Overall, we concluded that *Tt* Pah2 coordinates magnesium using Asp155, in addition to other canonical residues in the HAD superfamily. Given Asp155 is conserved in all lipin/Pah PAPs, this defines a common feature and suggests Asp155 plays a key role in catalysis.

To determine structures of additional catalytic states, we soaked Δhelix D148N apo crystals with magnesium and a variety of substrates and transition state analogs. Soaking of tungstate, but not other chemicals that mimic the transition state (e.g. vanadate), resulted in a positive peak adjacent to the magnesium ion in the Fo-Fc map. We thus modeled this density as a tungstate bound to the catalytic Asp146 **(Fig. 4C)** that mimics the phospho-Asp transition state intermediate generated after DAG is released from the active site in the first step of the mechanism **(Fig. 2)**.

Tungstate induced several notable changes in the active site. This included reorientation of the positively charged Lys244 sidechain towards the negatively charged tungstate **(Fig. 4C)**, which is a canonical interaction for HAD phosphatases^20,22^. Additionally, there was conformational change of the Arg269 sidechain that redirected the positively charged guanidinium group of Arg269 towards the center of the active site **(Fig. 4C)**. Arg269 is conserved in all experimentally characterized magnesium dependent PAPs including human and mouse lipins, and Sc Pah1, but is replaced by a Lys residue in some predicted, uncharacterized PAP enzymes (**Fig. 5B**). The high degree of conservation, along with the tungstate induced conformational change, suggested Arg269 may also play an important role in catalysis and represents another signature motif of lipin/Pah PAPs.

**Figure 5.**
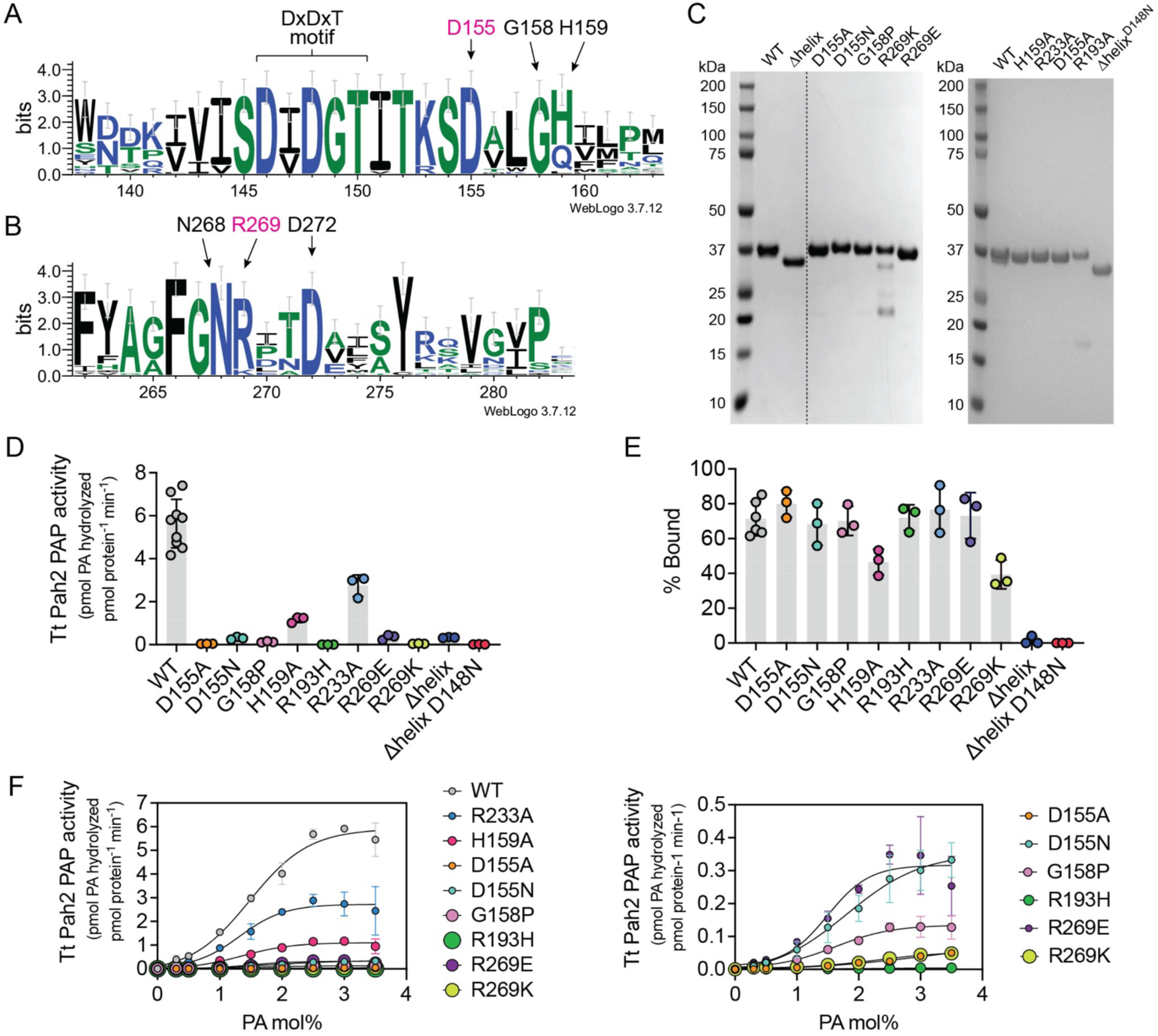
Conserved Arg and Asp residues are critical for *Tt* Pah2 PAP activity. **(A-B)** Sequence logos of (A) cap region 1 and (B) cap region 2 that highlights the conserved catalytic DxDxT motif in HAD phosphatases, and the conserved Asp and Arg residues (D155 and R269 in *Tt* Pah2) in lipin/Pah PAPs. **(C)** SDS-PAGE gel of purified *Tt* Pah2 proteins stained with Coomassie blue. **(D)** Effect of point mutations on the PAP activity of *Tt* Pah2. Δhelix denotes deletion of the membrane binding N-terminal helix. Error bars represent standard deviation (n = 3 for point mutants or n = 8 for wild-type *Tt* Pah2) of experiments performed in technical duplicates. **(E)** Quantification of liposome co-sedimentation for wild-type *Tt* Pah2 and point mutants expressed as % bound. Data are the means and SDs of three experiments. **(F)** Effect of point mutations on the PAP activity of *Tt* Pah2 as a function of the molar percent of NBD-PA (PA mol%) in Triton X-100 mixed micelles. The PA mol% was varied by maintaining a constant bulk concentration of NBD-PA (100 μM) and adjusting the amount of Triton X-100 detergent. Right panel shows a zoom up of point mutants with less than 5% activity relative to wild-type *Tt* Pah2.

### Asp155 and Arg269 are critical for activity

Next, we used point mutations to experimentally test the role of residues that our new structures suggested as important for PA binding and hydrolysis, including Asp155 and Arg269. Our goal was to identify residues involved in PA recognition that would be evolutionarily conserved to retain function. Thus, we chose to mutate residues that were both highly conserved, but also uniquely found in lipin/Pah PAPs. This strategy avoided altering residues absolutely conserved among all HAD phosphatases, as these residues would be necessary for catalytic function but were unlikely to directly be involved in PA binding.

In total, we generated eleven new point mutants of *Tt* Pah2. Four point mutants (D155R, G158A, R269H, and R269A) were either unstable and/or degraded, and could not be characterized. However, we were able to purify and characterize seven point mutants. This included two point mutants of Asp155 (D155N and D155A), two point mutants of Arg269 (R269K and R269E), two point mutants of highly conserved residues in cap region 1 (G158P and H159A), and one point mutant in cap region 2 (R233A). Arg233 was specifically chosen over other residues in cap region 2 because it was the only residue that was directed into the active site in any of our structures. For comparison, we included the R193H point mutant we previously characterized, and which represents the R725H point mutant found in lipin 1 deficient patients with rhabdomyolysis^11,19,30,34,39–41^.

Using fluorescent NBD-PA as a substrate solubilized in Triton X-100 mixed micelles, we compared the PAP activity of the points mutants to wild type (WT) *Tt* Pah2, the inactive R193H point mutant, and the Δhelix WT and D148N point mutants. All activity measurements were determined in the linear range of protein concentration, which varied depending on the relative activity of each point mutant. We found that only two point mutants, H159A and R233A, present in cap region 1 or 2 respectively, retained a significant amount (>20%) of PAP activity. The remaining point mutants resulted in less than 5% PAP activity compared to wild type *Tt* Pah2 (**Fig. 5D**). However, notably these point mutants did have measurable activity, unlike the R193H and the Δhelix D148N point mutants, which were completely inactive.

To control for the potential loss of membrane association, which would indirectly affect activity, we also analyzed membrane binding of all point mutants using a liposome co-sedimentation assay. As previously reported, the Δhelix constructs completely lost membrane association, whereas most point mutants had no effect on membrane binding **(Fig. 5E)**. The two exceptions were H159A and R269K that had an ∼50% reduction in membrane binding, which may account for some fraction of the loss of activity in these point mutants (**Fig. 5E**).

Next, we analyzed the activity of each point mutant as a function of mole percentage of PA in each detergent micelle (PA mol%). Wild type *Tt* Pah2 had a sigmoidal increase in PAP activity relative to increased PA mol%. The point mutants R233A, H159A, and G158P, while having reduced activity did not affect the shape of the curve. In contrast, the D155A and R269K point mutants had linear increases in activity with increasing PA activity, which suggested they had reduced PA affinity, in addition to a reduction in kcat (**Fig. 5F**). The effects of D155N were more modest suggesting the Asn residue may be able to sustain some minimal catalytic function. Surprisingly, the R269E point mutant had higher activity than the R269K point mutant, and a similar sigmoidal curve shape in comparison to WT. While we do not have an explanation for the ability of R269E to retain some function, given that D155 and R269 were the only residues that when mutated, affected the shape of the curve, we concluded that the Asp155 and Arg269 residues were most likely to be involved in PA recognition.

### Docking of PA suggests a mechanism for PA recognition by Arg269

To gain further insight into how lipin/Pah PAPs engage PA during catalysis and the residues involved in recognition, we performed flexible docking calculations using a water soluble, short chain PA and the D148N Δhelix magnesium bound crystal structure of *Tt* Pah2 (Structure 3). The D148N mutation was manually changed back to an Asp residue. The CHARMM force field^42^ and the CDOCKER^43^ method were using to perform docking calculations. A total of 1,030 flexible docking simulations were performed. Docking results generating structures with overall docking energies less than -50kJ/mol, distances shorter than 4.0 Å between the phosphate headgroup of PA and either the Mg^2+^ ion or the Oψ of Asp146, with PA in an orientation that would be deemed physiologically relevant, and with the phosphate headgroup located distal to the catalytic Asp and the fatty acid tails directed away from the active site, towards the membrane interface were considered favorable.

Analysis of the favorable docking poses revealed three major findings that all supported a role for Arg269 in PA recognition and Asp155 in catalysis (**Fig. 6A**). First, Arg269 consistently underwent a conformational change that moved the sidechain of Arg269 into the active site to make direct interactions with the sn2 glycerol backbone of PA (**Fig. 6B**). Second, Asp155 remained coordinated to the Mg^2+^ ion, but was found to alternate between two orientations that differed with respect to which oxygen atom of the carboxylate group bound magnesium. Lastly, Lys244 consistently underwent a conformational change that placed the amine group of the sidechain in position to interact with an oxygen of the phosphate headgroup of PA, similar to the tungstate bound crystal structure. This conformational change is observed in many HAD phosphatase structures, indicating that the enzyme was poised for catalysis in these docking structures. Several of the docking results that fit these criteria also had side chain orientations of amino acids in the active site nearly identical to that of the tungstate bound crystal structure (**Fig. 6C**), further validating the independent findings of the flexible docking.

**Figure 6.**
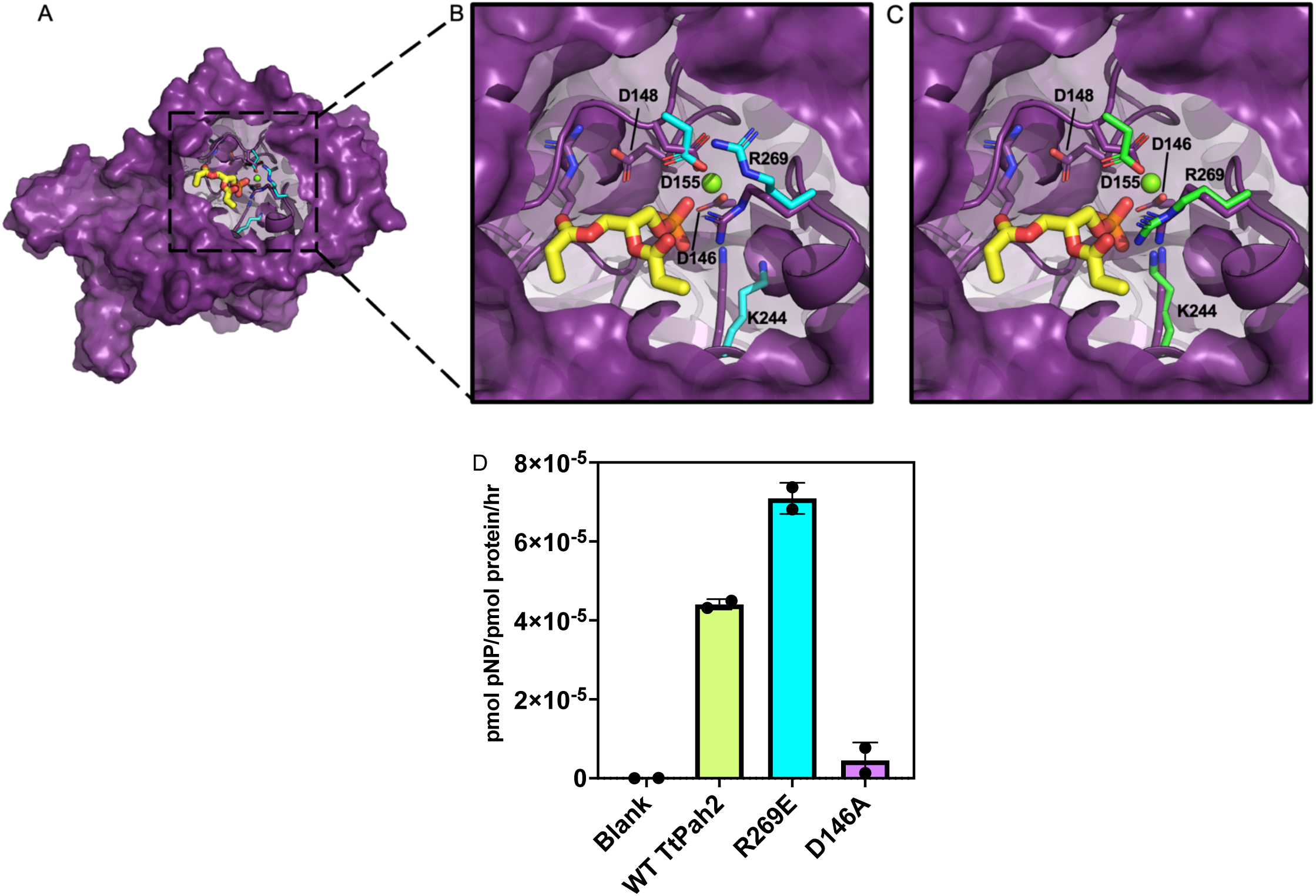
The role of Arg269 in recognition of the substrate phosphatidic acid. Docking results with favorable energies reveal conformational changes of Arg 269 side chain, similar to crystal structures. **(A)** Top-down view of the active site from a representative docking result (purple) with short chain phosphatidic acid (PA, yellow). **(B)** Inset: Overlay of active sites of a representative docking result (purple) and magnesium bound crystal structure (cyan), demonstrating reorientation of Arg269 and Lys244 when PA is present in the active site. **(C)** Inset: Overlay of active sites of a representative docking result (purple) with the magnesium and tungstate bound crystal structure (green), demonstrating side chain orientation similarity. The magnesium ion is represented as a green sphere and water soluble PA molecule docked in shown in yellow. **(D)** Detection and quantification of *Tt* Pah2 wild-type and mutant phosphatase activity against the substrate para-nitrophenol phosphate (pNPP).

### Arg269 is selectively required for PA catalysis

Our current structural and biochemical data supported a role for Arg269 in PA recognition. In particular, our docking analysis suggested Arg269 may engage the sn2 glycerol backbone of PA. To further characterize the role of Arg269, we compared the activity of wild-type *Tt* Pah2 with the R269E point mutant against the soluble substrate paranitrophenol-phosphate (pNPP). Using high concentrations of purified *Tt* Pah2 we were able to observe detectable pNPP hydrolysis, albeit at a low rate of ∼1 pNPP molecule hydrolyzed by every *Tt* Pah2 molecule per hour. A comparison of wild type *Tt* Pah2 with R269E *Tt* Pah2 observed that the R269E point mutant had no significant effect on *Tt* Pah2’s ability to hydrolyze pNPP (**Fig. 6D**). Thus, the detrimental effects of this point mutant on catalysis are limited to the endogenous substrate PA, consistent with a role of Arg269 in substrate recognition.

## DISCUSSION

The improved resolution of the structures presented above allowed us to establish a key role for Asp155 in Mg^2+^ coordination, and Arg269 in PA substrate specific recognition. Given their conservation, our findings support the role of these residues as a novel, signature motif of Mg^2+^-dependent lipin/Pah PAPs that distinguishes them from other HAD phosphatases. In addition, this study captures both global and local active site conformational changes of *Tt* Pah2. Overall, this provides insight into how the HAD phosphatase domain of lipin/Pah PAPs have evolved to act on their membrane embedded lipid substrate PA, and provides a framework that enables future experimental and computational studies to consider how membrane and lipid binding may influence the structural dynamics in lipin/Pah PAPs, especially the peptides present in cap regions 1 and 2.

The interactions observed between the sidechain guanidinium group of Arg269 with the sn2 glycerol backbone of PA in our flexible docking analysis provides one possible explanation for the preference of lipin/Pah PAPs for PA over the glycerol-3-phosphate pathway precursor lyso-PA, which lacks the sn2 acyl chain (**Fig. 1**). In fact, a role for Arg269 in catalysis and PA recognition was unexpected, as it is not part of the canonical catalytic residues of HAD phosphatases and resides outside of the canonical cap regions 1 or 2 that mediate substrate recognition in other HAD phosphatases. However, several independent lines of evidence point to a key role for Arg269 in *Tt* Pah2 and lipin/Pah PAPs in general. This includes the structural changes induced upon tungstate binding that reposition Arg269 over the active site, the effects on KM along with a reduction in kcat upon substitution of Arg269 to either a Glu or Lys residue, and the near universal conservation of Arg269 in both experimentally characterized and predicted lipin/Pah PAPs.

Previously, we postulated that *Tt* Pah2 would recognize its substrate solely via the loops in cap region 1 or cap region 2, consistent with Tt Pah2 being a HAD phosphatase with a type 0 cap. Studies of genetic evolution of this superfamily demonstrate that members have a similar Rossmann-fold at their core, and the caps evolved gain-of-function structural changes that are responsible for substrate recognition^20,31,32^. However, our data implicate Arg269 as being partially responsible for substrate recognition, although it lies neither in cap region 1 or 2. Notably, many other HAD phosphatases with cap type 0 act upon large macromolecular substrates, which is reasonable explanation for why they have do not have caps that occlude their active sites. *Tt* Pah2 acts upon a relatively small lipid, PA, which could explain differences in structural evolution and substrate recognition mechanisms from other type 0 HAD members. *Tt* Pah2 having a residue that plays a role in substrate recognition outside the cap regions, could in part explain how this HAD superfamily member acquired the unique biological function of acting on a membrane embedded substrate with high specificity, without having the active site occluded by a domain. This provides new insight of Mg^2+^-dependent lipin/Pah PAPs evolution as members of the HAD phosphatase superfamily.

Our work also implicates residues in cap regions 1 and 2 as important for *Tt* Pah2 PAP activity, with cap region 1 appearing most critical. Like the Mg^2+^-coordinating residue Asp155, Gly158 is also universally conserved in lipin/Pah PAPs and mutation to Proline greatly diminished activity. Unfortunately, the minimum alteration of Gly158 to Ala (G158A) resulted in protein degradation, which precluded further characterization. However, taken together, this raises the hypothesis that cap region 1 may undergo conformational change after either membrane or PA binding during catalysis.

Our prior structures provided a rational explanation for disease-associated point mutants in lipin PAPs, as *Tt* Pah2 conserves all of the native residues that are mutated in lipin associated pathologies. The new higher resolution structures presented here, give more confidence in the positioning and interactions for the native, unmutated residues **(Fig. 4)**, and also explain new disease-associated point mutants that have been identified in lipins. This includes D804N and D804H in lipin-1, which results in loss-of-function and rhabdomyolysis^33,34^. While we did not biochemically characterize the analogous point mutant in *Tt* Pah2, the Mg^2+^ bound structure reveals that the corresponding Asp residue (D272 in *Tt* Pah2) hydrogen bonds with a water molecule that occupies one of the Mg^2+^ coordination sites **(Fig. 4B,C)**. This provides a simple explanation for loss of activity.

In addition, the R736H/C (R193 in *Tt* Pah2) mutations have been reported in patients with lipin-2 loss-of-function causing Majeed syndrome^44,45^. Interestingly, the analogous mutation in lipin-1 R725H also results in loss-of-function, causing rhabdomyolysis. Our high-resolution structure **(Fig. 4D)** further confirms the importance of this Arg residue, demonstrating that it forms a salt bridge with the second Asp in the catalytic motif.

## METHODS

### Plasmids

The *Tetrahymena thermophila* Pah2 (*Tt* Pah2) gene encoding residues 1-321, which was previously codon optimized for E. coli expression and cloned into the pET28 plasmid with an N-terminal His-Tag, was used as a template to clone Tt Pah2 21-321 (Δhelix) and Tt Pah2 21-321 with a C-terminal His-Tag (Δhelix-His) into ppSUMO, which contained an N-terminal His-Tag followed by the ULP-1 cleavable SUMO protein. Point mutations of Tt Pah2 1-321 were made in pET28 vector by site directed mutagenesis. All sequences were verified by direct sequencing.

### Protein overexpression and purification

Plasmids were transformed into *E. coli* BL21 (DE3) RIPL cells (Agilent Technologies) for overexpression. Cells were grown at 37°C in Ultra-High Yield Flasks (Thompson Instrument Company) in Terrific Broth media to an optical density (OD) 600 nm of 1.5 and then cooled at 10°C for 2h. Protein expression was induced by addition of 100 μM isopropyl β-D-1-thiogalactopyranoside (IPTG) at 15°C overnight. Cell pellets were harvested by centrifugation at 4,000 RPM (3,320 x g) at 4°C for 20 min, flash frozen and stored at -80°C.

Frozen cell pellets were resuspended in either buffer A1 [50 mM Tris pH 8, 500 mM NaCl, 60 mM imidazole, 5% (v/v) glycerol, and 2 mM beta-mercaptoethanol (βME)] or buffer A2 [100 mM phosphate buffer, pH 7, 500 mM NaCl, 60 mM imidazole, 5% (v/v) glycerol, 0.5mM phenylmethanesulfonyl fluoride (PMSF) and 2 mM beta-mercaptoethanol (βME)] with addition of 1% Triton X-100, and lysed by sonication at 85% amplitude through 6-8 cycles of 2 sec on/off for 1 min. The D155A, D155N, G158P, R269K, and R269E cell pellets were lysed in buffer A2, while all other proteins were lysed in buffer A1. Proteins in a soluble fraction were collected by centrifugation at 26,000 RPM (81,770 x g) at 4°C for 1h. *Tt* Pah2 proteins were first purified by nickel-nitrilotriacetic acid (Ni-NTA) immobilized metal affinity chromatography (IMAC) using a 5mL HisTrap FastFlow column (GE Healthcare) and eluted in buffer A1 or A2 that contained an increased imidazole concentration of 300 mM. The *Tt* Pah2 proteins with an N-terminal SUMO fusion (Tt Pah2 21-321 (Δhelix) and Tt Pah2 21-321 with a C-terminal His-Tag (Δhelix-His) were then subjected to overnight incubation at 4°C with Ubl-specific protease 1 (ULP-1) for N-terminal 6x-His-SUMO fragment removal. All proteins with only an N-terminal His-Tag were directly applied to size exclusion chromatography. Next, proteins were further purified by size exclusion chromatography (SEC). Here, protein was passed through a 0.22 μm filter and applied to a Superdex 75 26/60 HiLoad column (GE Healthcare) equilibrated with SEC buffer [20 mM Tris pH 8, 150 mM NaCl, 5% glycerol (v/v), 10 mM βME, and 1 mM dithiothreitol (DTT)]. Protein fractions were concentrated by centrifugal ultrafiltration to ∼1-10 mg/mL, aliquoted, flash frozen, and stored at -80°C. Protein concentration was determined by UV absorbance spectroscopy using a NanoDrop (Thermo Scientific) and a calculated extinction coefficient at 280 nm.

### Crystallization and data collection

Purified *Tt* Pah2 ΔHelix D148N (21-321) protein at 3.82 mg/ml was initially screened for crystallization in 384 conditions using the JCSG core suites I-IV (Qiagen) in a sitting drop vapor diffusion format where the ratio of protein to mother solution was 1:1. This identified one promising condition: 0.1M CHES (pH 9.5) and 30% PEG 3000 for crystal growth. Optimization using a hanging drop vapor diffusion method and microseeding using a seed bead (Hampton research) generated multiple singular “X -like” crystals with 30% PEG 3000 and a pH range of 0.1 M CHES pH 8.5 - 9.5. The volume ratio of the protein to mother solution to undiluted seeds was 0.75 μL to 1 μL to 0.25 μL, respectively. All screening and crystal optimizations were performed at room temperature. Crystals were generated in 2-7 days. For data collection, crystals were incubated in a cryoprotectant solution plus a soaking agent and then flash-frozen in liquid nitrogen. Incubation times were 5 min to 1h. Soaking agent was 50 mM MgCl2 alone or with added 5 mM of one of the following: VnO4^3^, WO4^2-^, AlF4^-^, 6:0 PA, or G3P. The cryoprotectant solution consisted of mother liquor and 25% glycerol or ethylene glycol. The Tt Pah2 D155N was crystallized in a different condition consisting of 20% PEG 8000, 0.1 MES pH 6, and 0.2 M Ca Acetate. D155N crystals were cryoprotected using mother liquor with 25% glycerol. Native diffraction data were collected at the Advanced Photon Source (APS) 23 ID-D beamline at Argonne National Lab or at Brookhaven National Lab (BNL) NSLS-II FMX (17-ID-1) or AMX (17-ID-2) beamlines. All data were processed using xia2 DIALS in CCP4^46–48^.

### Structure determination and refinement

Phases for Structure 1 of *Tt* Pah2 ΔHelix-His were determined by molecular replacement (MR) carried out in PHENIX^49^, using Phaser^50^ with the previously reported structure of *Tt* Pah2 3.0 Å (PDB: 6TZZ) as a search model. Phases for Structure 2 were determined using MR using Structure 1 as a search model. Structures 3, 4, and 5 were determined using MR with Structure 2 as a search model. Manual model building in Coot^51^ with several rounds of refinement in Phenix produced the final model. The geometric quality of the refined model was evaluated with the structure validation tools in the Phenix suite. Data collection and refinement statistics are shown in Table 1. Structural figures were generated using the PyMOL 2.4 software.

### *Tt* Pah2 activity assay

*Tt* Pah2 activity was measured by fluorometric detection of NBD-PA undergoing hydrolysis to NBD-DAG^52^. The area of product was then converted to the moles of DAG produced based on a standard curve using known concentrations of NBD-DAG. Triton X-100 mixed micelles containing 14:0-12:0 NBD-PA (Avanti Polar Lipids, Catalog #810172) were prepared by drying NBD-PA in glass tubes under N2(g), resuspension in the appropriate amount of Triton X-100 (to determine the NBD-PA mol%), followed by multiple rounds of water bath sonication and vortexing. The protein was diluted in 2X activity assay buffer [40 mM Tris pH 7.5, 200 mM NaCl, 20 mM MgCl2, and 20 mM beta-mercaptoethanol (βME)]. 50 μL of NBD-PA (final concentration = 100 μM NBD-PA) Triton X-100 mixed micelles were incubated with 50 μL of protein for 30 min at 30 °C. The final reaction conditions contained: 20 mM Tris pH 7.5, 100 mM NaCl, 10 mM MgCl, and 10 mM beta-mercaptoethanol (βME). Each reaction was quenched with the addition of 300 μL of 1:1 CHCl3/MeOH (Sigma-Aldrich), followed by vortexing, and centrifugation at 2000 rpm (771 x g). The organic phase was collected, dried under N2(g), and resuspended with 100 μL of mobile phase B. Chromatographic separation was achieved utilizing an Agilent 1100 Series HPLC and mobile phase A and B eluents with a flow rate of 0.5 mL/min. Conditions were optimized using a Peek Scientific C-8 column (3 μm particle, 3.0 × 150 mm). Mobile phase A consisted of HPLC grade water and mobile phase B consisted of HPLC grade methanol. Both eluents were buffered by 0.2% formic acid (Fisher Chemical) and 1 mM mass spec grade ammonium formate (Sigma-Aldrich). Ratio of methanol/water during elution was 50:50. All enzyme assays were conducted in technical duplicates with a minimum of three experimental replicates. All reactions were linear with respect to time and protein concentration.

### Phosphatidic acid docking using the CDOCKER protocol

The 1.95Å *Tt* Pah2 crystal structure with a Mg^2+^ ion bound in the active site (Structure 3) was used as a starting structure (**Fig. 1D**). Structure 3 had a loop in cap region 1 with no density due to high heterogeneity and flexibility of these residues. The missing residues were graphically added back by editing the pdb file. Asn148 was also changed back to the Asp present in wild type *Tt* Pah2. This starting structure was then solvated and validated used the CHARMM input generator. The water-soluble PA substrate with C2 acyl chains was constructed using Avogadro^53^, to output a *mol2* formatted file. The *mol2* file was used as input for the cgenff program^54^, which generated the molecular mechanics force field parameters. The CDOCKER method^43^ and CHARMM version 41^42^ were used for the computational docking of the water-soluble PA substrate to the magnesium bound crystal structure of *Tt* Pah2 (**Fig. 1D**). The CDOCKER method begins with random rotations about each of the rotatable bonds of the substrate, followed by random translational and rotational motions of the substrate within the box defined by the user^43^. For these calculations, the cube was centered around the active site of *Tt* Pah2. The 20Å^3^ cube was centered around the magnesium ion resolved in the crystal structure and the catalytic residues, Asp146 and Asp148. The full molecular mechanics representation of the protein was then replaced with a *‘soft’* grid potential. The soft grid potential allowed the geometry to equilibrate, regardless of whether the initial geometry has strong repulsive potential between the enzyme and substrate. These optimizations were followed by simulated annealing molecular dynamics in the presence of the soft grids. This was followed by geometry optimization in the presence of the molecular mechanics potential energy function for the enzyme. A total of 1,030 individual docking calculations were performed for *Tt* Pah2 and the water-soluble PA substrate.

### pNPP Assay

A paranitrophenol phosphate (pNPP) master mix was prepared by the addition of 200 mM pNPP to assay buffer containing 20 mM Tris pH 8, 150 mM NaCl, 20 mM MgCl2, 20 mM BME. The reactions were carried out in a 96-well plate by mixing 50 μL of pNPP master mix with 20 μL of enzyme at 56 µM final protein concentration and 30 µL of assay buffer without BME. The absorbance at 405 nm was monitored with SpectraMax M2e Microplate Readers (Molecular Devices) at 30 s intervals at ambient temperature for 30 min. The moles of pNP product were quantified using a standard curve of product pNP created by dissolving pNP in the above assay buffer and measuring the absorbance at 405 nm of a 100 μL solution in a 96-well plate in triplicate.

## ACKNOWLEDGEMENTS

This work was supported by the National Institutes of Health grant R35GM128666 (to MVA), a Howard Hughes Medical Institute (HHMI) Gilliam Fellowship GT15815 (to FSW), an American Heart Association predoctoral fellowship 19PRE34450192 (to VIK), and a Sloan Research Fellowship (to MVA).

